# Automated Agnostic Designation of Pathogen Lineages

**DOI:** 10.1101/2023.02.03.527052

**Authors:** Jakob McBroome, Adriano de Bernardi Schneider, Cornelius Roemer, Michael T. Wolfinger, Angie S. Hinrichs, Aine Niamh O’Toole, Christopher Ruis, Yatish Turakhia, Andrew Rambaut, Russell Corbett-Detig

**Affiliations:** Department of Biomolecular Engineering, University of California Santa Cruz, Santa Cruz, CA, USA; Genomics Institute, University of California Santa Cruz, Santa Cruz, CA, USA; Biozentrum, University of Basel, Basel, Switzerland; Swiss Institute of Bioinformatics, Basel, Switzerland; Department of Theoretical Chemistry, University of Vienna, 1090 Vienna, Austria; Research Group Bioinformatics and Computational Biology, Faculty of Computer Science, University of Vienna, 1090 Vienna, Austria; RNA Forecast e.U., 1100 Vienna, Austria; Institute of Ecology and Evolution, University of Edinburgh, Edinburgh, UK; Molecular Immunity Unit, University of Cambridge Department of Medicine, MRC-Laboratory of Molecular Biology, Cambridge, UK; Department of Veterinary Medicine, University of Cambridge, Cambridge, UK; Cambridge Centre for AI in Medicine, University of Cambridge, Cambridge, UK; Department of Electrical and Computer Engineering, University of California San Diego, San Diego, CA, USA

## Abstract

Pathogen nomenclature systems are a key component of effective communication and collaboration for researchers and public health workers. Since February 2021, the Pango nomenclature for SARS-CoV-2 has been sustained by crowdsourced lineage proposals as new isolates were added to a growing global dataset. This approach to dynamic lineage designation is dependent on a large and active epidemiological community identifying and curating each new lineage. This is vulnerable to time-critical delays as well as regional and personal bias. To address these issues, we developed a simple heuristic approach that divides a phylogenetic tree into lineages based on shared ancestral genotypes. We additionally provide a framework that automatically prioritizes the lineages by growth rate and association with key mutations or locations, extensible to any pathogen. Our implementation is efficient on extremely large phylogenetic trees and produces similar results to existing Pango lineage designations when applied to SARS-CoV-2. This method offers a simple, automated and consistent approach to pathogen nomenclature that can assist researchers in developing and maintaining phylogeny-based classifications in the face of ever increasing genomic datasets.

## Introduction

Pathogen nomenclature systems, or the designations of groups below the level of species, are important for facilitating effective research, treatment, and communication about diseases. Despite the universal importance and long history of nomenclature systems for pathogens, there remains a plurality of approaches to apply to new emerging pathogens. These nomenclature systems are generally based on some combination of three elements: phenotype, genotype, and geography. Phenotype-based systems are often predicated on vulnerability to antibiotics (Collins et al. 1982) or serology (Lancefield 1933); pathogens with serology based nomenclature systems include Salmonella spp. (Brenner et al. 2000), dengue viruses (Cuypers et al. 2018; Simmonds et al. 2017), and Streptococcus spp. (Facklam 2022; Lancefield 1933). Geography-based classification systems may be appropriate for pathogens where the primary reservoir is in non-human species, such as Chikungunya virus (CHIKV) (de Bernadi Schneider et al. 2019) and the Zaire Ebola viruses (Kuhn et al. 2014). Finally, genotype-based nomenclature divides a species-wide phylogeny into statistically well-supported, mutually exclusive taxa generally referred to as “lineages” or “clades”. These groups can be defined as clusters of samples below a genetic diversity threshold or as the descendents of an inferred common ancestor on a single phylogeny. Genotype-based classification has become increasingly common in application to viruses such as RSV (Ramaekers et al. 2020), dengue (Cuypers et al. 2018) and influenza viruses (Anderson et al. 2016).

The Pango system is a genotype-based dynamic lineage nomenclature for SARS-CoV-2 characterized by the manual designation of new lineages from a global phylogenetic tree (Rambaut et al. 2020). Pango lineages are hierarchical and comprehensive, including hundreds of nested designations for any subgroup of viruses that may be of concern. The Pango system has provided initial names used for all Variants of Concern (VOCs), including B.1.1.7 (Alpha) and B.1.1.529 (Omicron), and defined the serial replacement of Omicron lineages through time (BA.1, BA.2, BA.5). Pango has played a critical role in facilitating effective tracking of and communication about emerging SARS-CoV-2 strains over the course of the pandemic. The maintenance and continued efficacy of this system is therefore of critical importance for public health.

Currently, Pango relies on manual curation and designation, including the crowdsourcing of lineage proposals on a public forum (https://github.com/cov-lineages/pango-designation). More than 2500 SARS-CoV-2 variants have been named under the Pango system as of January 2023, an average of more than two new lineages per day since the beginning of the pandemic. The trained human eye is excellent at distinguishing new lineages of interest from groups of low-quality or contaminated isolates, but the Pango group’s resources are becoming strained as the volume of data increases and public investment decreases. Furthermore, crowdsourced proposals are vulnerable to delays as well as regional and personal bias, as individual researchers have differing opinions on the importance of various mutations and are more or less likely to search for clades from specific parts of the world. A more objective metric to evaluate candidates for lineage designation could help to reduce this bias and streamline the lineage proposal and review process.

Additionally, it is uniquely challenging to define lineages for SARS-CoV-2, due primarily to low genetic diversity and massive volumes of data. Other diseases often have lineages with common ancestors inferred to have been circulating several years to decades in the past, with well-defined characterizing genetic changes. These stable nomenclatures rarely need active updates, being defined with respect to a single phylogeny that remains largely unchanged. In SARS-CoV-2, however, a single mutation may be all that defines a new epidemiologically distinct lineage (O’Toole et al. 2021). The volume of SARS-CoV-2 sequencing data is also orders of magnitude greater than other pathogens, straining existing systems (Hodcroft et al. 2021). The SARS-COV-2 phylogeny itself is regularly updated (McBroome et al 2021), necessitating further review and updates to any lineage nomenclature. Pango is therefore a dynamic nomenclature, needing constant maintenance as new epidemiological lineages emerge. This makes it substantially more burdensome to maintain through manual surveillance. A computable metric to automatically identify and sort lineage candidates would assist curators in building and maintaining the Pango system. Moreover, as pathogen genome sequencing expands in other species, similar difficulties in rapid lineage designation will become commonplace.

We propose a simple heuristic approach for the definition and expansion of genotype-based dynamic nomenclature systems. Our method is rooted in information theory, optimizing for the representation of sample-level haplotype information. It requires only a phylogeny with branch lengths scaled to genetic distance and a set of user parameters. It is efficient in application to extremely large phylogenies and produces a comprehensive hierarchy of genetically distinct lineages. Our lineage system is flexible and can be effectively weighted in any number of ways, allowing epidemiologists and researchers working with any pathogen to prioritize critical elements for lineage definition and tracking efforts. Importantly, it can expand a preexisting lineage system, making adoption of this approach for the maintenance and expansion of existing nomenclature straightforward. We, in collaboration with the Pango designation team, have implemented this system as a new input for the existing Pango lineage designation infrastructure (https://github.com/jmcbroome/autolin). This approach can easily be generalized, and as sequencing technology becomes more widely applied, our scalable and generalizable method can be applied to produce and maintain nomenclature systems for novel and extant pathogens.

## Results

### The Genotype Representation Index

A nomenclature system can be likened to a language, where additional words, analogous to lineages, are defined for common, unique concepts to reduce the average number of words per sentence. Along these lines, an effective nomenclature summarizes a complex phylogeny into useful, distinct categories to facilitate effective analysis and communication. The lineage hierarchy is generally defined with respect to a specific rooted phylogeny, where a number of specific ancestral nodes are designated as lineage roots (Supplementary Figure 1). Higher-level lineages are divided hierarchically into finer sublineages. Individual samples, represented as tips of the tree, are members of every lineage that is rooted in its inferred ancestry. To automate the definition of this hierarchy, we need an objective measure of distinctiveness or importance that can be computed for individual nodes across the tree. Once we have a measure of lineage efficacy, we can iteratively construct a nomenclature by identifying high-value nodes and designating them as lineage roots. These lineages can then be presented to an end user, or directly incorporated into an expanding nomenclature.

To this end, we define the following index, hereafter referred to as the “genotype representation index” (GRI) (Figure 1).

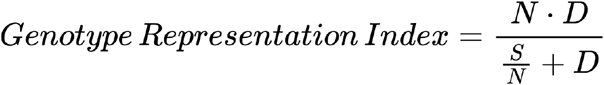

**Figure 1:**
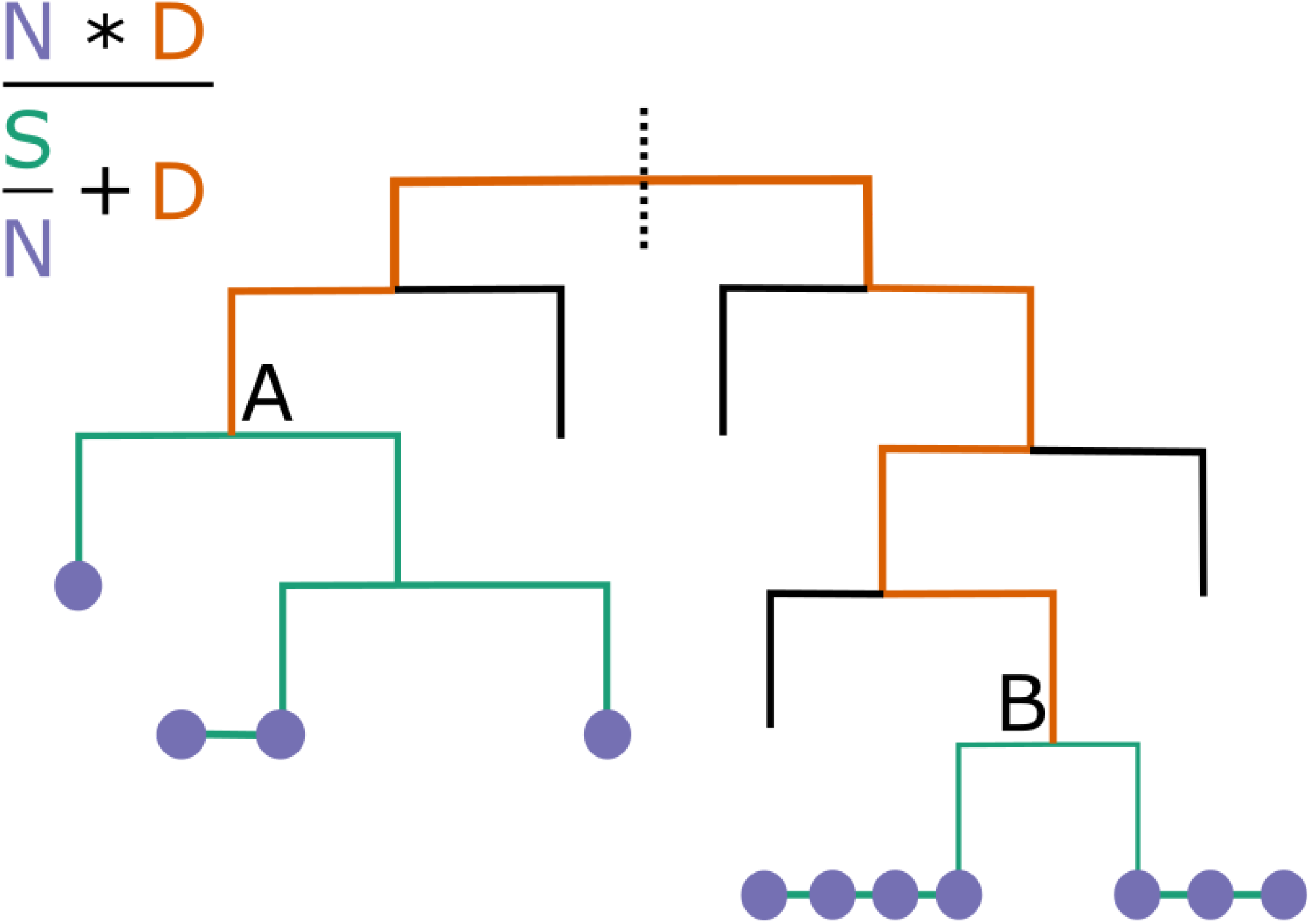
Computation of GRI. This cartoon depicts two candidate nodes for lineage definition. The path to the root is in orange, paths to each descendent in teal, and descendants represented by lilac circles. Lilac circles on the same horizontal line are genetically identical. All vertical lines represent a single mutation, hence node A has a GRI of ~2.1, and node B has a much larger GRI of ~5.6. The B node is more desirable as a lineage label because it represents a distinct and large group of low-diversity samples. Such a group of samples, with many shared mutations, is more likely to be an epidemiologically distinct category.

Here, N is the number of descendent tips from the node, D is the total branch length from the node to the root of the tree or parent lineage, and S is the total branch length from the node to each descendent tip. For example, for a mutation-annotated tree (Turakhia et al 2021), such as those used for SARS-CoV-2, the branch length (D and S) is in units of total mutations across the genome. However, the GRI can be computed on any rooted tree topology, as long as branch lengths are scaled by genetic distance. The GRI is high for internal nodes where descendent samples are genetically similar to one another and the node itself is genetically distinct from the rest of the phylogeny, desirable qualities for lineage designation (Rambaut et al. 2020).

To define a lineage system based on the GRI, we apply a greedy maximization algorithm. The GRI is computed for each node on the tree and the node with the highest value is chosen as a new lineage root. Additional mutually exclusive lineages are defined by disregarding all samples covered by an existing lineage label and recomputing the GRI for all remaining samples and their ancestors. Additional hierarchical lineages are defined similarly by only considering samples descended from an existing “parent” lineage root. This process is repeated until a desired number of lineage labels have been defined or all available nodes fail to pass thresholds for designation. This iterative approach is not guaranteed to find the highest overall GRI lineage configuration among many possible combinations of lineages, but it scales well to millions of samples and a rapid pace of lineage updates.

The GRI is rooted in information theory; the node with the highest value will convey the most information on average if given a lineage label. Additional details on the motivating theory can be found in the Methods section.

### Adjustments to the Genotype Representation Index

In practice, pathogen lineage nomenclature systems are generally designed for purposes beyond summarization of a phylogenetic tree. Lineages often carry connotations of distinct phenotypic behavior, such as serological types, immune evasion, transmissibility, and other metrics. Some parts of the genome may contribute more than others to these phenotypes. For example, some spike protein changes are known to alter immune escape in SARS-CoV-2 (Greaney et al. 2022). Information about parts of the genome associated with important phenotypes is inherently more valuable and more worth representing in our lineage system. Accordingly, we may want to weight our GRI calculation by giving additional value to these mutations when computing distances. Conversely, we may want to disregard parts of the genotype that are not informative for phenotypic behavior or that are not readily interpretable, such as repetitive noncoding sequences or sites prone to recurrent errors.

The GRI, while based on genotype representation, can be flexibly altered to focus on the representation of important elements. The original Pango rules for the definition of SARS-CoV-2 lineages have requirements around evidence for international transmission and changes to proteins to designate lineages that are more likely to be epidemiologically important (Rambaut et al. 2020). By using these weighting schemes for GRI, we can automatically propose lineage designations of high epidemiological import. This allows researchers to develop fully-informed and highly applicable nomenclature systems.

### Example Lineages

To demonstrate the utility of our approach, we applied our method to the global public phylogeny as of 2022-12-11 from http://hgdownload.soe.ucsc.edu/goldenPath/wuhCor1/UShER_SARS-CoV-2/ (McBroome et al. 2021). We generated 187 new lineage designations using the default configuration parameters, which only considers samples collected in the preceding 8 weeks (Supplementary File 1). 24 of these lineages were actively sampled in December 2022 as of 2022-12-11. These active designations were highly dispersed in size, with a mean size of 82 samples and a median of 45 samples. The full report for the active designations is available in Supplementary Table 1.

We fit an exponential growth model to each active lineage (see Methods) (Supplementary Table 1; Figure 2) and obtained a 95% confidence interval estimate of the rate of exponential growth. The average confidence interval for the exponential growth interval was relatively large (0.07, 0.49), due primarily to the effects of limited sample sizes. 16 of the 24 lineages had a positive lower interval bound, which is evidence for active spread in the countries they are present in. The width of the interval is dependent on the data available; while the average estimate for our lineages is +/− 0.2, estimates for lineages with at least 50 total collected samples had a much narrower average value of +/− 0.07. All model confidence intervals are reported in Supplementary Table 1. All code for fitting and reproducing these results is available at https://github.com/jmcbroome/lineage-manuscript.

**Figure 2:**
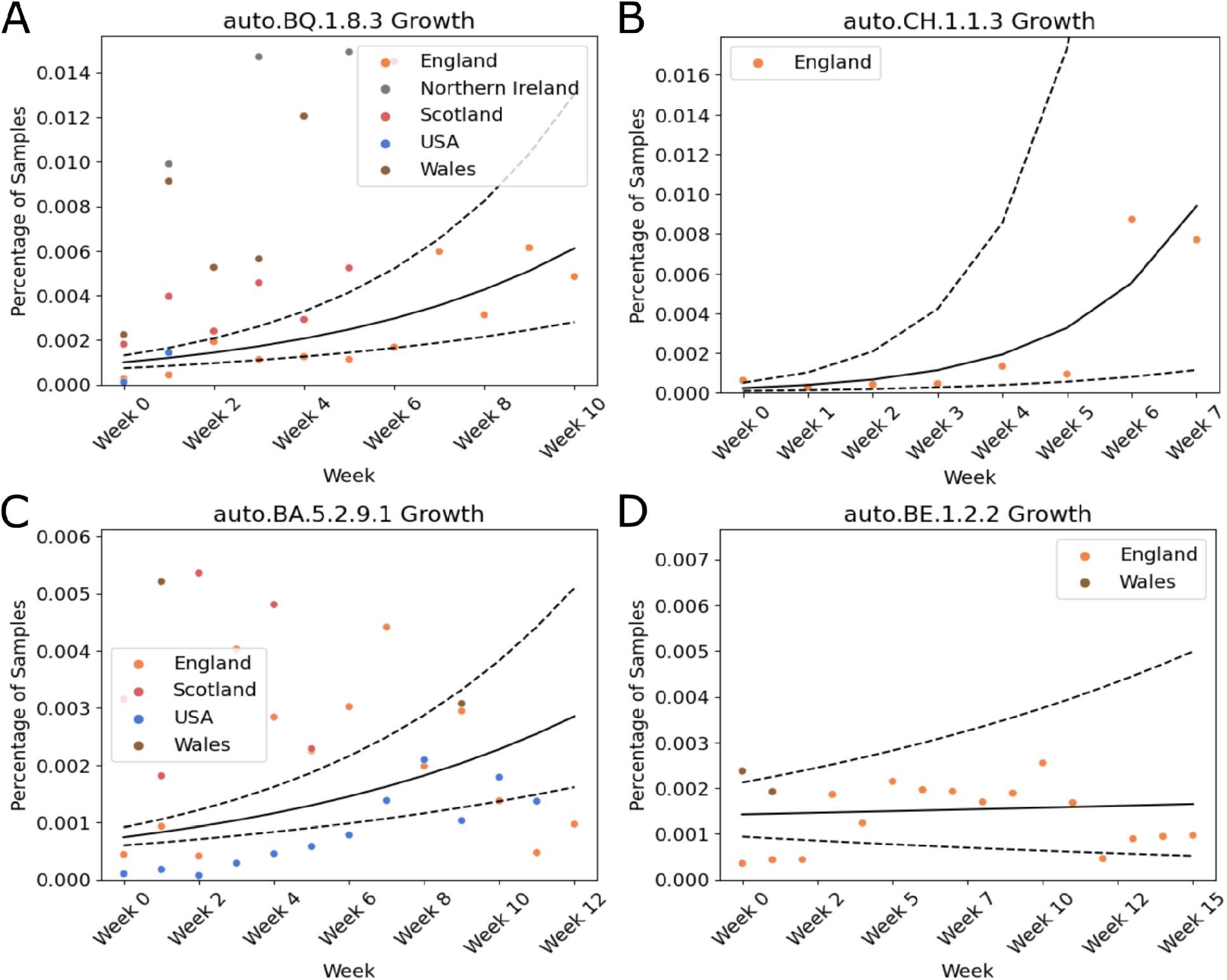
Exponential Growth Modeling. The above four plots describe some of the lineage annotations produced by our method based on the public SARS-CoV-2 data. The black line is the median estimated growth trajectory, while the dotted lines represent the lower and upper bounds of the 95% confidence interval of the growth trajectory. The x-axis is represented in weeks since first detection among each country.

Figure 2 displays a small selection of lineages and model fits in further detail. The naming schema matches the Pango naming schema, with the addition of an “auto” prefix denoting that the lineage in question was created by our approach and not manually designated by the Pango team. “auto.CH.1.1.3”, while exclusive to England, exhibits a very rapid expansion in latter weeks that drive a very high, if wide, estimate of growth. “auto.BQ.1.8.3” and “auto.BA.5.2.9.1” are more international, but less consistent; the latter appears to grow consistently in the United States, but fluctuates to a much greater degree in England. Finally, “auto.BE.1.2.2.” is an example of a low-priority designation, with no strong evidence of positive growth. Altogether, our models are capable of capturing a diverse set of lineage trajectories and rapidly and effectively identifying lineages undergoing exponential expansion.

We have collaborated with the Pango team to incorporate our approach into the existing SARS-CoV-2 lineage designation infrastructure. Statistics such as lineage size, associated mutations, and geographic localization are computed and reported as a part of a pull request to the curated Pango repository. Our update includes links to external data exploration sources such as cov-spectrum (Chen et al. 2022) and taxonium (Sanderson 2022; Kramer et al. 2022), as well as programmatic generation of all files requisite for the incorporation of the new designations. All code for this procedure can be found at https://github.com/jmcbroome/autolin.

### Application to Other Pathogens

The GRI approach can be applied outside of updating dynamic nomenclature like Pango. We evaluated its potential for automatically producing stable lineage systems based on more traditional phylogenies with Chikungunya virus (CHIKV, geographic nomenclature) and Venezuelan equine encephalitis virus complex (VEE, serology based nomenclature) using the currently available nextstrain builds (CHIKV Nextstrain build 5.1 (https://nextstrain.org/groups/ViennaRNA/CHIKVnext) and VEE Nextstrain build 2.1 (https://nextstrain.org/groups/ViennaRNA/VEEnext). Overall the CHIKV geographic nomenclature aligns with the automated lineage designations at its base level (ARI=0.69, p=0.018), with further breaking down of the tree in certain regions such as the Indian Ocean Lineage (Figure 3). On the other hand, VEE’s serology based nomenclature is paraphyletic and does not represent phylogenetic lineages or clades (Forrester et al. 2017, de Bernardi Schneider & Wolfinger 2023). We elected to present two levels of annotation, reflecting the distinction between VEE viruses generally and the Venezuelan Equine Encephalitis Virus (VEEV) and its subtypes. VEEV itself is successfully identified from the VEE complex by our lineage approach at the first level of annotation (ARI=0.9, p=0.0003). However, our method was unable to reliably recapitulate VEEV serotypes at the second level of annotation (ARI=0.28, p=0.25, Figure 4), due largely to the paraphyletic nature of VEEV’s serotype-based nomenclature. Regardless, these two examples show how this method can generate de novo lineage classification of pathogens, independent of context and consistent with human intuition. Our implementation for the generation of lineage labels for arbitrary pathogens can be found at https://github.com/jmcbroome/automated-lineage-json. It is provided as both a command line interface tool and as an online Streamlit app, accessible at https://jmcbroome-automated-lineage-json-streamlit-app-3adskh.streamlit.app/. This demonstrates that the GRI approach can be applied in a pathogen-agnostic fashion to produce nomenclature consistent with existing systems.

**Figure 3:**
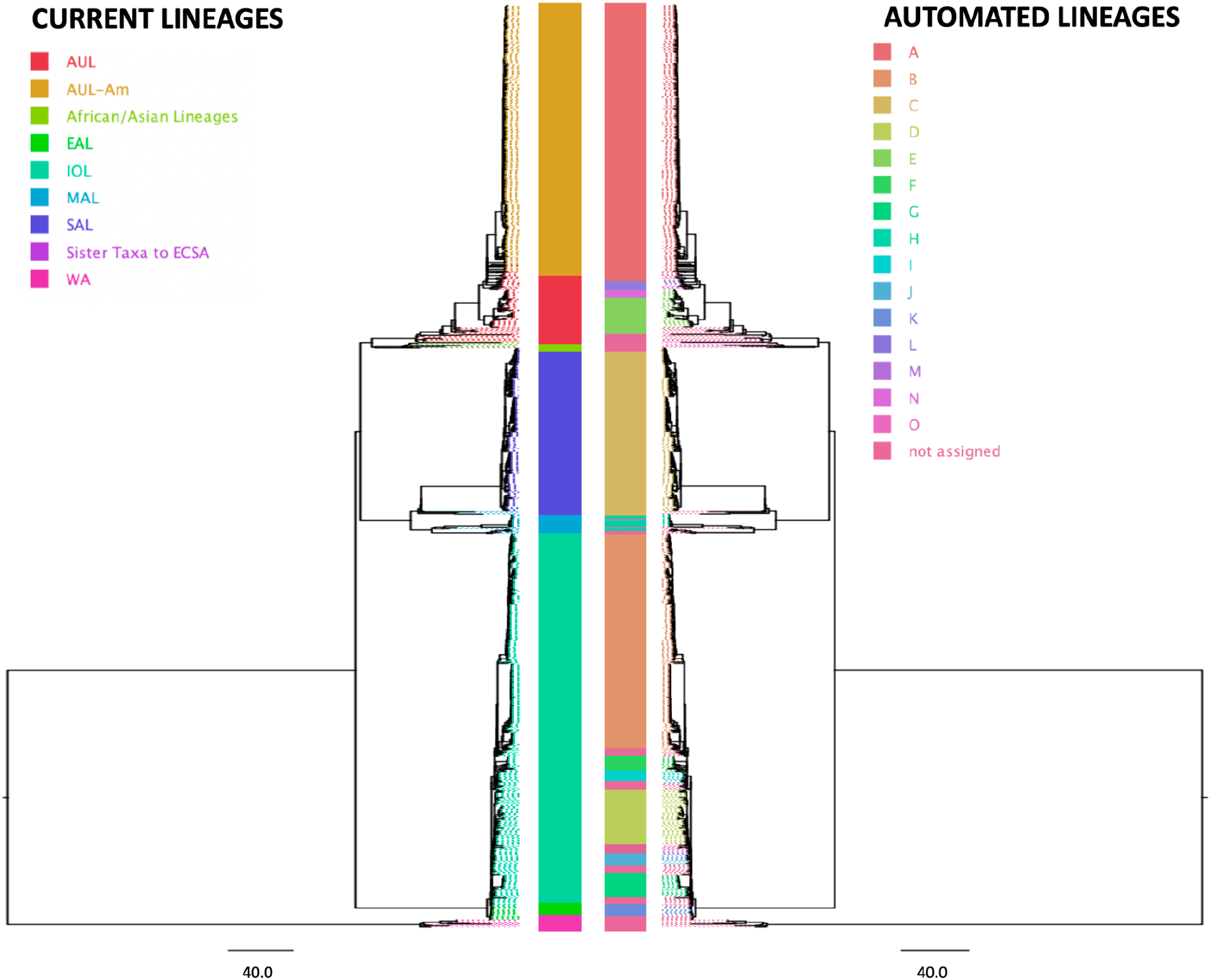
Comparison of the geography lineage designation (left tree) with automated lineage designation (right tree) of Chikungunya virus, based on a tree previously generated by the Augur pipeline (Huddleston et al 2021) and visualized on FigTree v.1.4.4.

**Figure 4:**
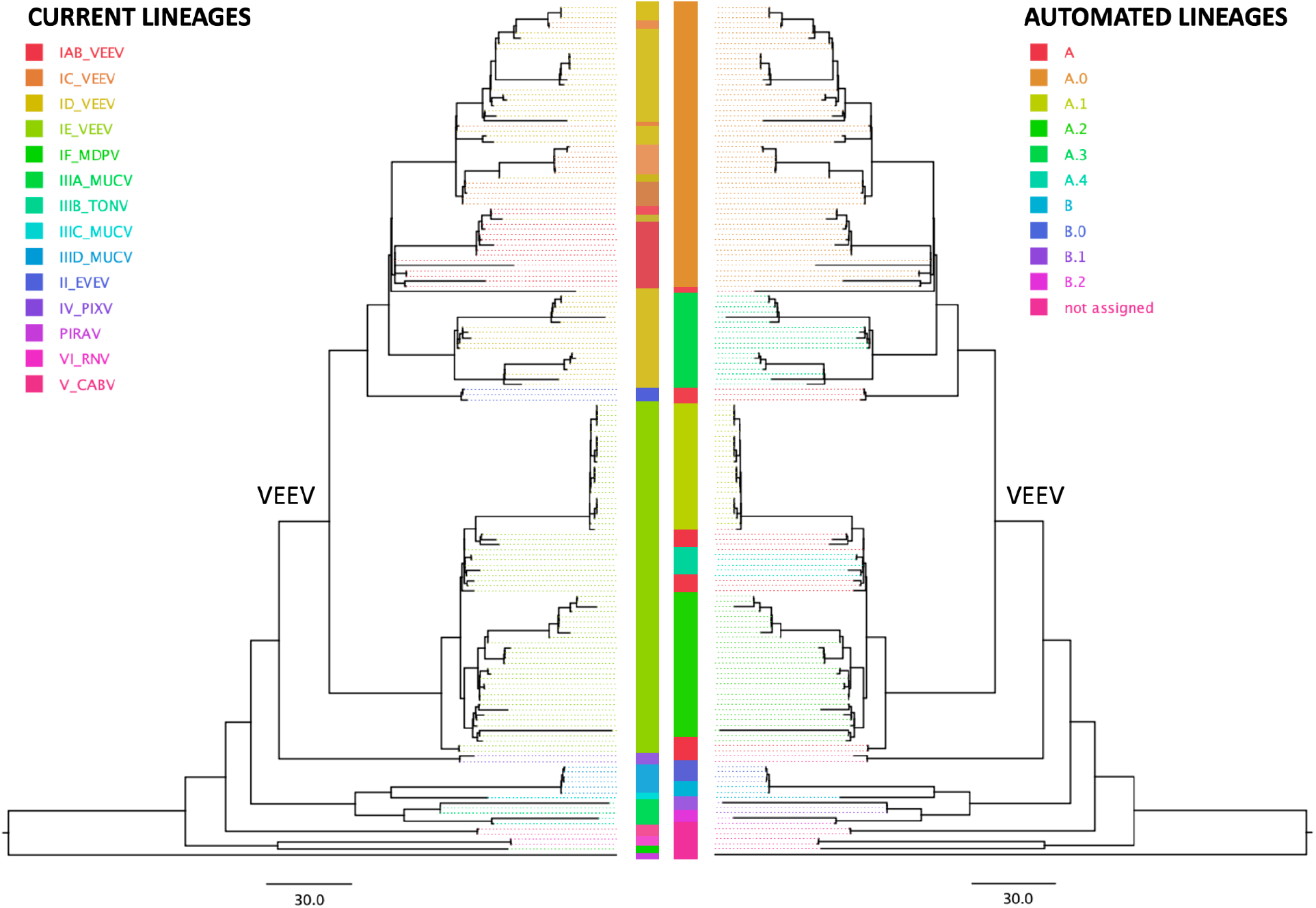
Comparison of the serology subtype designation (left tree) with automated lineage designation (right tree) of the Venezuelan equine encephalitis virus complex (VEE), based on a tree previously generated by the Augur pipeline (Huddleston et al 2021) and visualized on FigTree v. 1.4.4. According to the current nomenclature, VEE encompasses Everglades virus (EVEV), Mucambo virus (MUCV), Tonate virus (TONV), Pixuna virus (PIXV), Cabassou virus (CABV), Rio Negro virus (RNV), Mosso das Pedras virus (MDPV), Pirahy virus (PIRAV) and the Venezuelan equine encephalitis virus (VEEV). The VEEV clade is labeled in the tree.

## Discussion

We have presented a new index-based method for the generation of genotype-based nomenclature, capable of both expanding existing nomenclature systems and generating novel nomenclature on understudied or emerging pathogens. While designed for the demands of the SARS-CoV-2 pandemic, this approach can be easily applied to any rooted tree with branch lengths scaled by genetic distance, such as those created and distributed by Nextstrain (Hadfield et al. 2018). Our implementation is efficient and includes several parameters to adjust the behavior of the metric. These include prioritizing the labeling of specific mutations or specific tips and only considering mutations effects on specific proteins.

Nonetheless, our approach does exhibit a few potential issues, shared with many lineage nomenclatures. First, it is defined with respect to a specific phylogeny. This can be problematic when attempting to maintain lineages over time, as new data is collected and the phylogeny is updated. Phylogenetic inference is naturally uncertain, and optimization of an existing phylogeny may alter lineage relationships or invalidate identified lineages. The dual system of comprehensive, automatically designated clades and a curated subset of epidemiologically distinct lineages is largely robust against these issues, as manual curators may be more likely to spot problematic clades and prevent them from being designated as full lineages. In rare cases, lineages may need to be retracted or redefined, as is the case for current Pango lineages when new data suggest alternative relationships than the one originally used for lineage designation.

Second, SARS-CoV-2 recombines at low rates (Jackson et al. 2021; Turakhia et al. 2022). The apparently long branches which occur on the phylogeny as a result will often be picked up by this method as a new lineage annotation, but the ancestry of that lineage annotation cannot be accurately represented by a single tree topology. Incorporating new samples or optimizing the phylogeny may relocate it from the acceptor to the donor parent lineage, vice versa, or elsewhere on the phylogeny altogether. In this scenario, a recombinant lineage may have to be retracted or renamed. Alternatively, lineages identified as recombinants can receive special designation names and we have previously developed methods for comprehensively identifying recombinant lineages within SARS-CoV-2 phylogenies (Turakhia et al. 2022) that may facilitate this effort.

SARS-CoV-2 is likely to become an endemic pathogen, similar to the *influenza virus* (Otto et al. 2021). Accordingly, there is likely to be a long-term pattern of replacement of existing strains, demanding ongoing designation of new lineages for effective monitoring of pathogen diversity (Rambaut et al. 2020). Investing into infrastructure to reduce manual curation will lead to long-term consistency and effectiveness of designation. Additionally, it is likely that automated approaches will be faster than many human-based systems, thereby promoting stability of public and scientific discourse by labeling potentially important lineages before they are widespread and contributing to major epidemiological patterns worldwide. The results we present here may serve for consistent, immediate SARS-CoV-2 lineage designation for years to come.

Overall, this approach for lineage designation is generic, flexible, and applicable to future datasets with unclear nomenclature or expansive phylogenies. At the time of publication it is somewhat limited in practical application, with SARS-CoV-2 being the only pathogen with the scale of data to strain direct manual curation. However, global pathogen sequencing is on the rise, and generalized concepts for the creation of new nomenclature or expansion of existing systems, as we exemplified with CHIKV and VEE, will be critical for future public health challenges. There is currently no universal definition of nomenclature below the species level (ICTV Code), leading to widespread debate and confusion between epidemiological analyses. This method can serve to unify and streamline the definition and maintenance of future nomenclature systems.

## Acknowledgments

This research was funded in part by the Austrian Science Fund (FWF) project I 6440-N to MTW. This research was funded in part by CDC award BAA 200-2021-11554 to RBC. We gratefully acknowledge Olyver Pybus for contributing to early discussions on the topic. We also would like to thank all the volunteers who have worked on finding new Pango lineages.

## Methods

### Information Theoretical Underpinnings

A lineage system can be formulated as a sender/receiver information scenario. The sender possesses the full phylogenetic tree and a lineage system L, while the receiver possesses only the lineage system L and the associated mutation paths that define each lineage. For initial simplicity, we assume the lineage system consists of a single label, applied to branch B. Let the receiver be interested in the full ancestry of an arbitrary sample S. S may or may not be a member of a lineage L. If it is, the receiver already has all ancestry information associated with that lineage L for the sample S. How much additional information is required to specify the full ancestry of sample S?

A single site’s state can be represented in a finite number of bits; 2 bits to represent the state and 15 bits to represent the location, for SARS-CoV-2. Therefore, the full ancestry path of a given branch can be represented in a finite number of bits, proportional to the number of mutations separating it from the root.

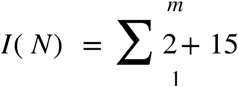

Therefore, the additional information required to specify the ancestry of sample S, given a lineage system with a label at branch B, is

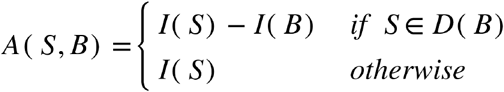

Where D(B) is the set of descendant samples from labeled branch B. This extends to lineage systems with multiple branches B by checking membership of each outermost branch label B. Given that the receiver may be interested in any arbitrary sample S, we choose to place our label at branch B such that the additional information required to specify the full ancestry of S is, on average, minimized.

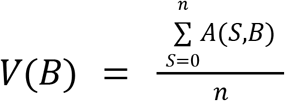

We choose B such that V(B) is minimized. To accomplish this, we need to maximize both the number of descendants of the branch we chose and the representation by that branch of those samples. The genotype representation index (GRI) is the mean of I(B) for all samples S descended from B, times the number of samples descended from B. We use the multiplied mean value of I(B) instead of the sum both for computational ease and to reduce the impact of outlier leaves with excessively long path lengths. The higher the GRI of branch B, the lower the A(S,B) of the system from that branch. As A(S,B) is reduced, so is V(B). Choosing the branch with the highest GRI as our lineage branch B will yield a system with a low V(B) and high informational efficiency.

Extending this approach to additional lineages is straightforward. “Serial” or non-overlapping lineages, where

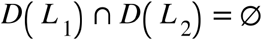

Can be assigned by repeating the minimization procedure while disregarding all samples that are a member of existing lineages. This can be repeated until some minimum percentage of samples are contained within some set D(L).

“Hierarchical” or nested lineages, where

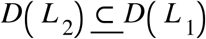

Can be assigned by treating L_1_ as the root of the tree, with ancestry information conveyed with respect to it. There are no other types of lineage relationship, as a rooted phylogenetic tree is a directed acyclic graph and lineages are always defined as a monophyletic clade. It is not possible for two clades to partially overlap when they are defined by internal nodes on a fixed phylogenetic tree.

If an arbitrary number of lineages are allowed to be labeled, eventually every internal node may be labeled as a unique lineage- the receiver will therefore already possess the exact and complete phylogeny, and additional information will never be necessary. However, this defeats the point of the lineage system, which is intended as an effective summary of the phylogeny that captures most relevant information without costing as much to store and manage.

We should consider that a lineage ought to include more than a mere handful of samples, even if those samples have relatively long unique paths. Many lineage systems require a minimum number of samples to be represented by that label. We therefore define a minimum *m*; we subtract the weighted mean information represented by a theoretical set of *m* samples with the same path length distribution from the true information distribution for the node. If the net information represented is negative, then we reject this node as a candidate for a new lineage definition.

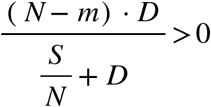

Essentially, we require that *N* > *m*, where *m* is a user selected parameter, in order to define a new lineage. Setting this to X will produce only proposed lineages that convey some information about at least X leaves.

Similarly, we can set a minimum distinguishing distance from the subtree root/parent lineage. Often lineage designation systems require some number of unique distinguishing mutations for a new sublineage. We therefore define

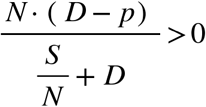

When *p* < *D*, this value is negative and we reject this candidate node. Setting this to two, for example, will produce only lineages that convey at least two unique mutations distinct from the parent lineage or tree root. Combining both of these filters, we reject nodes where either or both of these inequalities are not passed. Together, this allows automatic proposals to fulfill standard conditions required by lineage nomenclature review groups.

### Additional Parameters

Our pipeline implementation includes a substantial set of configurable parameters. These include minimum lineage size and minimum distinction, as outlined above. We also can simply threshold on the GRI itself, ignoring marginal designations that contain relatively little additional information.

Notably, we can additionally incorporate arbitrary sample-level weighting. This allows our lineage system to prioritize effective representation of high-interest samples. R(S), below, is a function representing the “importance” of sample S. This might be high for a sample S from an under sequenced region, or lower for a sample S from a heavily sequenced time or place.

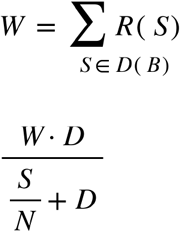

Samples from regions that contribute a small percentage of all samples will have substantially higher weights than ones from regions that contribute a large percentage of sequences, though all samples will have a weight greater than 1 under this schema. This is just one potential weighting schema for handling geographic sequencing bias, and the user can define any schema and set weights on a per-sample basis.

Similar concepts can apply to computing path lengths- we may consider only part of the haplotype, or assign additional weight to specific mutations of interest that we want our lineage system to prioritize representing. We provide options for the user to select genes of interest for representation, as well as the ability to ignore mutations that do not change amino acid content of proteins and represent coding haplotypes only.

We also provide arbitrary weighting schema for mutations of interest, similar to samples. As an example, we provide a parameter that heavily weights mutations that are predicted to increase vaccine escape (Greaney et al. 2022). This parameter multiplies the escape weight value estimated by the Bloom lab calculator by the user’s parameter and adding 1. In this schema, mutations that are not predicted to contribute to immune escape have a weight of 1, while mutations that do contribute have a weight greater than 1 that is proportional to the strength of escape conferred. The resulting lineage system is more likely to include designations that have a change in immune escape. This is just one possible schema and the user can define weights on a per-mutation basis in our implementation.

All parameters and configuration information used in the production of these results can be found in Supplementary File 1.

### Bayesian Growth Model

To identify high-priority lineages after designation, we fit a geographically stratified exponential growth model to each proposed lineage using Markov Chain Monte Carlo (MCMC) implemented in the package PyMC3. Bayesian methods of this type are appropriate for inference with small, noisy datasets, as the uncertainty in the model is directly quantified. To summarize, we construct a posterior distribution of exponential growth coefficient scores filtered through a binomial sequencing model. Lineage proposals which have a high, low-variance posterior distribution of growth are likely rapidly expanding and are of high priority for labeling.

For lineage L in country C, we model the true percentage P as increasing in an approximately exponential fashion. This is appropriate for newly emerging lineages that consist of a small percentage of total cases in any country where they are found but are successfully spreading. Each data point consists of the total number of samples from lineage L found in a specific country during a specific week. We assume that the inherent exponential growth coefficient for L is shared across all countries in which it is found and combine all datapoints across countries and times for each lineage. The first week that any sample from lineage L was found in country C is treated as the initial timepoint (t=0) for data from that country.

We do not directly observe the true percentage of cases P that are of lineage L. Instead, some number N of all cases are sequenced, and we observe some number X of these samples to be lineage L. As the number of cases is much larger than the number of samples, we can model this process as a binomial sampling procedure with N trials and a probability of success being the true percentage P.

Our Bayesian model combines both this sampling procedure and the exponential growth model to yield a posterior distribution of growth values which can explain the behavior of lineage L. Often these distributions are wide, due to sparse sampling and noise over few datapoints. Additionally, some lineages may not fit an exponential growth model at all, due to being outcompeted by newly introduced lineages or simple epidemiological noise, leading to highly variable estimates of growth. Accordingly, we compute the 0.025 and 0.975 quantiles (95% CI) for this distribution for each lineage L and sort the output by the lower quantile. Lineages with a large positive value for the lower quantile will reliably resemble a high exponential growth model and are more likely to be of epidemiological concern.

All code for our modeling and reporting process can be found at https://github.com/jmcbroome/lineage-manuscript and https://github.com/jmcbroome/autolin.

### Method validation - Other Pathogens

To validate that this method can be applied to pathogens other than SARS-CoV-2, we selected two nextstrain instances for Chikungunya virus and the Venezuelan Equine Encephalitis complex viruses, which are currently classified based on their geography and serology, respectively. We applied our generalized implementation (https://github.com/jmcbroome/automated-lineage-json) under default settings for the Auspice JSON files of each virus (CHIKV Nextstrain build 5.1 available at https://nextstrain.org/groups/ViennaRNA/CHIKVnext (doi:10.5281/zenodo.7514289) and VEE Nextstrain build 2.1 available at https://nextstrain.org/groups/ViennaRNA/VEEnext (doi:10.5281/zenodo.7524848)) to obtain lineage assignments. These Nextstrain JSON were generated by the Augur pipeline (nextstrain-augur v19.1.0, treetime v 0.9.4, iqtree v2.2.0). We then downloaded the nexus file with annotations from the new JSON file from Nextstrain and visualized and compared the annotations using FigTree v.1.4.4. Tree figure comparisons were made by extracting them in pdf format as displayed in FigTree, mirrored and aligned on a photo editing software. Taxon labels were colored according to the lineage assignment and were replaced with bars representing the color of the lineage for best visualization.

We compared the automated lineage assignments with the previous nomenclature using the Adjusted Rand Index (ARI). We randomly selected nodes in the amount of the number of categories found for each annotation to create a distribution of random ARI’s to evaluate the robustness of the method. By selecting random nodes within the tree and taking their descendants to construct our null comparisons, we account for natural correlation from the tree structure, while the Adjusted Rand Index itself accounts for variations in group sizes. We then compute the percentile of the true Adjusted Rand Index of our lineage proposals against the existing nomenclature from the permuted null distribution, yielding the reported p-values. All code for this can be found at https://github.com/jmcbroome/lineage-manuscript.

## Works Cited

Anderson, T. K. et al. A Phylogeny-Based Global Nomenclature System and Automated Annotation Tool for H1 Hemagglutinin Genes from Swine Influenza A Viruses. mSphere 1, e00275–16 (2016).

Brenner, F. W., Villar, R. G., Angulo, F. J., Tauxe, R. & Swaminathan, B. Salmonella Nomenclature. Journal of Clinical Microbiology 38, 2465–2467 (2000).

Chen, C. et al. CoV-Spectrum: analysis of globally shared SARS-CoV-2 data to identify and characterize new variants. Bioinformatics 38, 1735–1737 (2022).

Collins, C. H., Yates, M. D. & Grange, J. M. Subdivision of Mycobacterium tuberculosis into five variants for epidemiological purposes: methods and nomenclature. Epidemiology & Infection 89, 235–242 (1982).

Cuypers, L. et al. Time to Harmonize Dengue Nomenclature and Classification. Viruses 10, 569 (2018).

de Bernardi Schneider, A. et al. Updated Phylogeny of Chikungunya Virus Suggests Lineage-Specific RNA Architecture. Viruses 11, 798 (2019).

Wolfinger, M.T., and Schneider, A. de B. (2023). ViennaRNA/VEEnext: VEEnext v2.1. Zenodo. 10.5281/zenodo.7524848.

Facklam, R. What Happened to the Streptococci: Overview of Taxonomic and Nomenclature Changes. Clinical Microbiology Reviews 15, 613–630 (2002).

Forrester, Naomi L., et al. “Evolution and spread of Venezuelan equine encephalitis complex alphavirus in the Americas.” PLoS neglected tropical diseases 11.8 (2017): e0005693.

Greaney, A. J., Starr, T. N. & Bloom, J. D. An antibody-escape estimator for mutations to the SARS-CoV-2 receptor-binding domain. Virus Evolution 8, veac021 (2022).

Hadfield, J. et al. Nextstrain: real-time tracking of pathogen evolution. Bioinformatics 34, 4121–4123 (2018).

Hodcroft, E. B. et al. Want to track pandemic variants faster? Fix the bioinformatics bottleneck. Nature 591, 30–33 (2021).

Huddleston, J., Hadfield, J., Sibley, T.R., Lee, J., Fay, K., Ilcisin, M., Harkins, E., Bedford, T., Neher, R.A., and Hodcroft, E.B. (2021). Augur: a bioinformatics toolkit for phylogenetic analyses of human pathogens. Journal of Open Source Software 6, 2906. 10.21105/joss.02906.

ICTV Code: the International Code of Virus Classification and Nomenclature. International Committee on Taxonomy of Viruses (ICTV) https://talk.ictvonline.org/information/w/ictv-information/383/ictv-code/ (2018).

Jackson, B. et al. Generation and transmission of interlineage recombinants in the SARS-CoV-2 pandemic. Cell 184, 5179–5188.e8 (2021).

Kauffmann, F. The bacteriology of Enterobacteriaceae. Collected studies of the author and his co-workers. (1966).

Kramer, A. M., Sanderson, T. & Corbett-Detig, R. Treenome Browser: co-visualization of enormous phylogenies and millions of genomes. Bioinformatics 39, btac772 (2023).

Kuhn, J. H. et al. Nomenclature-and Database-Compatible Names for the Two Ebola Virus Variants that Emerged in Guinea and the Democratic Republic of the Congo in 2014. Viruses 6, 4760–4799 (2014).

Lancefield, R. C. A Serological Differentiation of Human and Other Groups of Hemolytic *Streptococci*. J Exp Med 57, 571–595 (1933).

McBroome, J. et al. A Daily-Updated Database and Tools for Comprehensive SARS-CoV-2 Mutation-Annotated Trees. Molecular Biology and Evolution (2021) doi:10.1093/molbev/msab264.

O’Toole, Á. et al. Assignment of epidemiological lineages in an emerging pandemic using the pangolin tool. Virus Evolution 7, veab064 (2021).

Otto, S. P. et al. The origins and potential future of SARS-CoV-2 variants of concern in the evolving COVID-19 pandemic. Current Biology 31, R918–R929 (2021).

Ramaekers, K. et al. Towards a unified classification for human respiratory syncytial virus genotypes. Virus Evol 6, veaa052 (2020).

Rambaut, A. et al. A dynamic nomenclature proposal for SARS-CoV-2 lineages to assist genomic epidemiology. Nat Microbiol 5, 1403–1407 (2020).

Salimi, V. et al. Proposal for Human Respiratory Syncytial Virus Nomenclature below the Species Level. Emerg Infect Dis 27, 1–9 (2021).

Salvatier, J., Wieckiâ, T. V. & Fonnesbeck, C. PyMC3: Python probabilistic programming framework. Astrophysics Source Code Library ascl:1610.016 (2016).

Sanderson, T. Taxonium, a web-based tool for exploring large phylogenetic trees. eLife 11, e82392 (2022).

Simmonds, P. et al. A proposed system for the nomenclature of hepatitis C viral genotypes. Hepatology 19, 1321–1324 (1994).

Simmonds, P. et al. ICTV Virus Taxonomy Profile: Flaviviridae. J Gen Virol 98, 2–3 (2017).

Spicher, T., Delitz, M., Schneider, A. de B. & Wolfinger, M. T. Dynamic Molecular Epidemiology Reveals Lineage-Associated Single-Nucleotide Variants That Alter RNA Structure in Chikungunya Virus. Genes (Basel) 12, 239 (2021).

Turakhia, Y. et al. Pandemic-scale phylogenomics reveals the SARS-CoV-2 recombination landscape. Nature 609, 994–997 (2022).

