## Supplementary Information for "Automated Agnostic Designation of Pathogen Lineages"

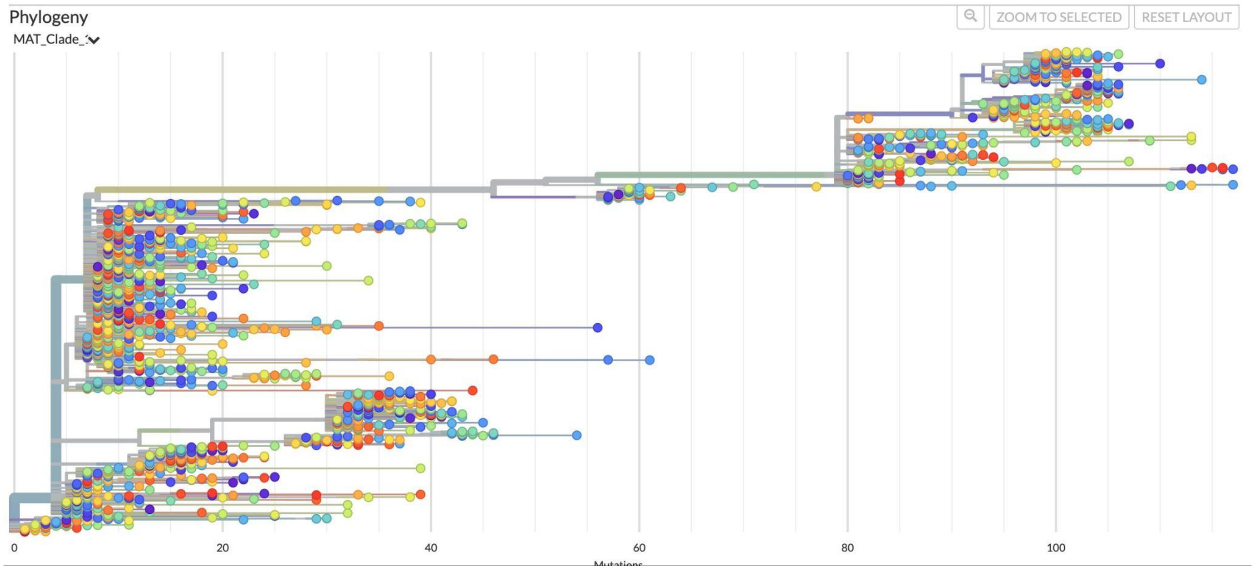


**Supplementary Figure 1: Pango Lineage Hierarchy.** This figure displays the hierarchical and serial relationships among the defined Pango lineages as of 2022-12-11.
An interactive version of this figure can be found at https://nextstrain.org/fetch/raw.githubusercontent.com/jmcbroome/lineage-manuscript/main/public-2022-12-11.backbone.json/?c=MAT_Clade_0&tl=MAT_Clade_0

**Supplementary Table 1: Output Report for 24 New Lineage Designations.** This table includes basic statistics for 24 new lineage designations actively sampled in December 2022.

**Supplementary File 1: Configuration for Output Report.** This file contains the configuration used to produce the lineages from the 2022-12-11 global public SARS-CoV-2 phylogeny in YAML format.
