## Supplementary File 1 for "Automated Agnostic Designation of Pathogen Lineages"

escape_data: "./SARS2_RBD_Ab_escape_maps/processed_data/escape_data.csv"
reference_genome: "./reference/NC_045512v2.fa"
reference_gtf: "./reference/ncbiGenes.noORF1a.gtf"
genbank: "./reference/hu1.gb"
python: python #path or name for a specific Python interpreter to use.
lineage_params:
  floor: 3 #minimum score value. Set higher to filter smaller and less valuable proposals. Set to 0 to report any and all sublineage proposals that pass other filters.
  annotation: B.1.1.529 #consider only this lineage and its descendents for sublineage proposal. Set to None to consider all lineages. 
  missense: True #consider only mutations that lead to amino acid changes as part of the haplotype. 
  gene: None #consider only mutations within the indicated gene as part of the haplotype. Set to None to use the whole genome.
  minsamples: 10 #minimum sample weight. Default weighting scheme is 1 per sample (more for underrepresented countries when country_weighting is used).
  distinction: 1 #minimum mutation weight. Default weighting scheme is 1 per mutation (more for mutations associated with immune escape if escape_weighting is used).
  recursive: False #allow proposal of additional sublineages from newly proposed lineages. Can break links for recursively proposed lineage issue output.
  cutoff: 0.99 #stop adding new serial (exclusive) lineage annotations when this percentage of samples is labeled.
  earliest_date: '2022-10-15' #ignore all samples before this date. Set to 2019-12-01 to examine the entire pandemic.
  weight_params:
    escape_weighting: 0 #Higher values means mutations associated with immune escape by the Bloom lab DMS are given higher weight. Set to 0 to disable escape weighting. 
    country_weighting: 10 #Higher values means samples from underrepresented countries count more. Set to 0 to disable country weighting. 
reporting_params:
  sort_by: "proposed_sublineage_score" #sort the results by this column.
  number: 20 #report the top X lineages in issues.
  prefix: "proposed_" #set a prefix for markdown and associated file output.
  samples_named: 15 #name up to this many samples within the text of the report, including the oldest and youngest samples. 
  local_only: True #set to True to print markdown issues to local files instead of attempting to post them to the repository issues board.
request_params:
  designation_repo: "./auto-pango-designation" #path to a local clone of https://github.com/cov-lineages/pango-designation or one of its forks. Required for pull request generation.
  local_only: True #update local files and print the text for a pull request body to a local markdown file instead of opening a branch and pull request on the targeted repository.
  auto_merge: False #immediately merge the generated pull request, if your api_key has permissions.
  representative_number: 1000 #add up to this many lineage representative samples per lineage to lineages.csv.
  valid_samples: "None" #use samples from this file when expanding lineages.csv. Set to "None" to use any samples from the tree.
  countries: 1 #set a minimum number of countries a lineage must exist in to be included in the request.
  maximum: 25 #set a maximum number of lineages to include in a pull request. The top N are sorted by stratified growth.
  active_since: '2022-12-01' #only include lineage proposals that were actively sampled since the indicated date. Set to "None" to report all.
  samples_different: 5 #block sublineage proposals that are less than this many samples different from the parent lineage. Set this higher to block proposals that constitute the vast majority of the parent lineage. Set to 0 to disable this filter. 
  growth_model: #parameters to adjust the behavior of the inferred growth model.
    use_model: True #set to True to infer the model, applying the parameters below, and add to the results table.
    draws: 1000 #set the drawing parameter for the sampler. https://www.pymc.io/projects/docs/en/stable/api/generated/pymc.sample.html
    tune: 1500 #set the tuning parameter for the sampler.
    target_accept: 0.9 #set the target acceptance parameter for the sampler.
    min_country_weeks: 2 #require this many total data points, as combinations of countries and weeks. 
    max_proportion_considered: .1 #ignore datapoints at a higher proportion than this for being unlikely to behave exponentially or exhibiting other issues.
